# A dominant role of TGFβ in regulating T-cell size and physiology

**DOI:** 10.1101/2024.06.02.597009

**Authors:** Inbal Eizenberg-Magar, Jacob Rimer, Stav Miller, Yehezqel Elyahu, Michal Mark, Ziv Porat, Shlomit Reich-Zeliger, Alon Monsonego, Yaron E Antebi, Nir Friedman

**Affiliations:** Department of Molecular Genetics, Weizmann Institute of Science, Rehovot 7610001, Israel; Department of Immunology and Regenerative Biology, Weizmann Institute of Science, Rehovot 7610001, Israel; Department of Life Sciences Core Facilities, Weizmann Institute of Science, Rehovot 7610001, Israel; The Shraga Segal Department of Microbiology, Immunology and Genetics, Faculty of Health Sciences, Ben-Gurion University of the Negev, Beer-Sheva 84105, Israel; Zlotowski Center for Neuroscience, and Regenerative Medicine and Stem Cell Research Center, Ben-Gurion University of the Negev, Beer-Sheva 84105, Israel; National Institute for Biotechnology in the Negev, Ben-Gurion University of the Negev, Beer-Sheva 84105, Israel; Department of Systems Immunology, Weizmann Institute of Science, Rehovot 7610001, Israel

**Keywords:** TGFβ, cell size, CD4^+^ T cells, regulation, cytokines, cellular processes

## Abstract

During an immune response, cells are simultaneously exposed to multiple cytokine signals that collectively determine their phenotype. Transforming growth factor β (TGFβ) is a pleiotropic cytokine acting as a key regulator of T-cell differentiation with activating and suppressive effects on their immune function. Here, we systematically analyze the cellular responses of CD4^+^ T cells to TGFβ across diverse cytokine environments in the presence or absence of TGFβ. We found that TGFβ had a profound dominant effect independent of the presence of other cytokines, modulating the expression of more than 4,000 genes. In the presence of TGFβ, cells exhibit lower expression of translation-related and apoptosis-related genes, accompanied by increased survival of activated T cells. Notably, cells cultured in the presence of TGFβ were smaller in size while preserving their proliferative ability. Accordingly, we identified a dense network of transcription factors that were modulated by TGFβ, suggesting a core gene set connecting TGFβ signaling to the regulation of T-cell size. We found N-Myc to be at the center of this network, and we directly show that TGFβ regulates its gene expression level, protein level, and nuclear localization. Our work provides a system to study cell size control and demonstrate the profound effect of TGFβ in the modulation and regulation of T-cell properties, expanding its role beyond guiding their phenotype.

**Significance Statement:** TGFβ is a key determinant of CD4^+^ T-cell differentiation; however, understanding its effect on additional aspects of T-cell state is lacking. Here, we systematically studied the role of TGFβ in regulating T-cell physiology. Exposing cells to diverse combinations of cytokines enabled us to distill the core effect of TGFβ. We found TGFβ to have a profound effect on multiple cellular processes critical to T-cell function. Significantly, TGFβ induced smaller T-cells both in vitro and in vivo, suggesting that TGFβ could skew the population towards tissue infiltration and residency. Furthermore, TGFβ can be used to fine-tune T-cell size, providing a system for studying cell size control. Overall, our findings demonstrate the profound effect of TGFβ in the regulation of T-cell physiology.

## Introduction

CD4^+^ T cells, central mediators of adaptive immunity, are activated upon T-cell receptor (TCR) recognition of antigenic peptides in MHC-II context and respond by changing a large set of cellular properties. The latter include abrupt changes in cell size, rapid proliferation accompanied by cell death, changes in metabolism (1, 2), and differentiation into specialized lineages with distinct immunological functions (3). These cellular processes depend on external signals that CD4^+^ T cells sense during their activation, such as co-stimulatory and co-inhibitory molecules, and cytokines that the cells encounter during antigen priming (3).

Transforming growth factor β (TGFβ) is a pleiotropic cytokine that influences many types of cells in different stages of development. Of note, TGFβ can have opposing effects on cells, depending on the cellular context (4–8). For example, TGFβ promotes tissue growth and morphogenesis and induces vasculogenesis and angiogenesis during embryonic development. In contrast, in mature tissues, the majority of cell types respond to TGFβ with growth arrest or cell death (9). In the immune system, TGFβ plays a central role in the regulation of inflammation. It significantly impacts T cells throughout different stages of their development, starting from thymic development, maintaining homeostasis in the periphery, affecting tolerance to self-antigens, and modulating proliferation, differentiation, and survival (5, 7, 10, 11). In combination with other cytokines, TGFβ drives the differentiation of CD4^+^ T cells towards several T helper (Th) cell subsets. The effect of TGFβ on T cells can be either stimulatory or inhibitory, depending on the state of the cells and the presence of additional regulatory signals. For example, in combination with IL-2, TGFβ drives the differentiation of cells towards the anti-inflammatory Treg lineage, while together with IL-6 or IL-4 it drives the formation of the pro-inflammatory lineages, Th17 or Th9, respectively (12–16). In a previous work, we conducted a systemic analysis of cytokine combinations and found that TGFβ had the strongest effect on CD4^+^ T cell fate amongst tested cytokines (17). While TGFβ is extensively studied in the context of CD4^+^ T-cell differentiation, a comprehensive understanding of its effect is still lacking.

Here, we used a combination of experimental techniques to comprehensively explore the effects of TGFβ on key cellular processes of CD4^+^ T cells after their activation. We applied TGFβ in combinations with different cytokines to focus on the functions of TGFβ that are independent of any specific signaling milieu and decouple its effect from that of other cytokines. Using gene expression analysis, we revealed that TGFβ affects a large set of genes, which are involved in a number of central cellular processes. Combining cell cytometry and imaging techniques, we found that TGFβ influences CD4^+^ T-cell proliferation and supports survival. Moreover, we show that cytokine combinations that include TGFβ consistently generated smaller T cells. Based on existing literature, we have curated a dense protein interaction network that underlies TGFβ regulation of cell size, and identified N-Myc as a possible key mediator of this effect. Together, our findings reveal that TGFβ plays a crucial role in regulating T-cell characteristics during immune responses and provide new insights on signal hierarchy during T-cell differentiation and fate determination.

## Results

### TGFβ in combination with other cytokines substantially modifies the gene expression profile of CD4^+^ T cells

To study the effects of TGFβ on CD4^+^ T cells, we started by examining genome-wide changes in gene expression profiles after exposure to TGFβ. Naïve CD4^+^ T cells were isolated from spleens of C57BL/6 mice and activated in vitro via the TCR and costimulation, using plate-bound αCD3 and αCD28 antibodies, respectively, together with mixtures of additional cytokines. Specifically, we used eight cytokine combinations: IL2, IL4, IL6, and IL12 with and without TGFβ. To distill the core effect of TGFβ from that of other cytokines, we focused on responses that consistently depend on TGFβ, regardless of its partners in the cytokine mix. As an additional control, we compared the effect of IL6 in combination with IL4 or IL12, as we have previously found IL-6 to have a modifying effect on CD4^+^ T cells (17) (Fig. 1A). Following 96 hours of culture, the cells were harvested, and genome-wide mRNA expression was measured using RNA sequencing.

**Figure 1.**
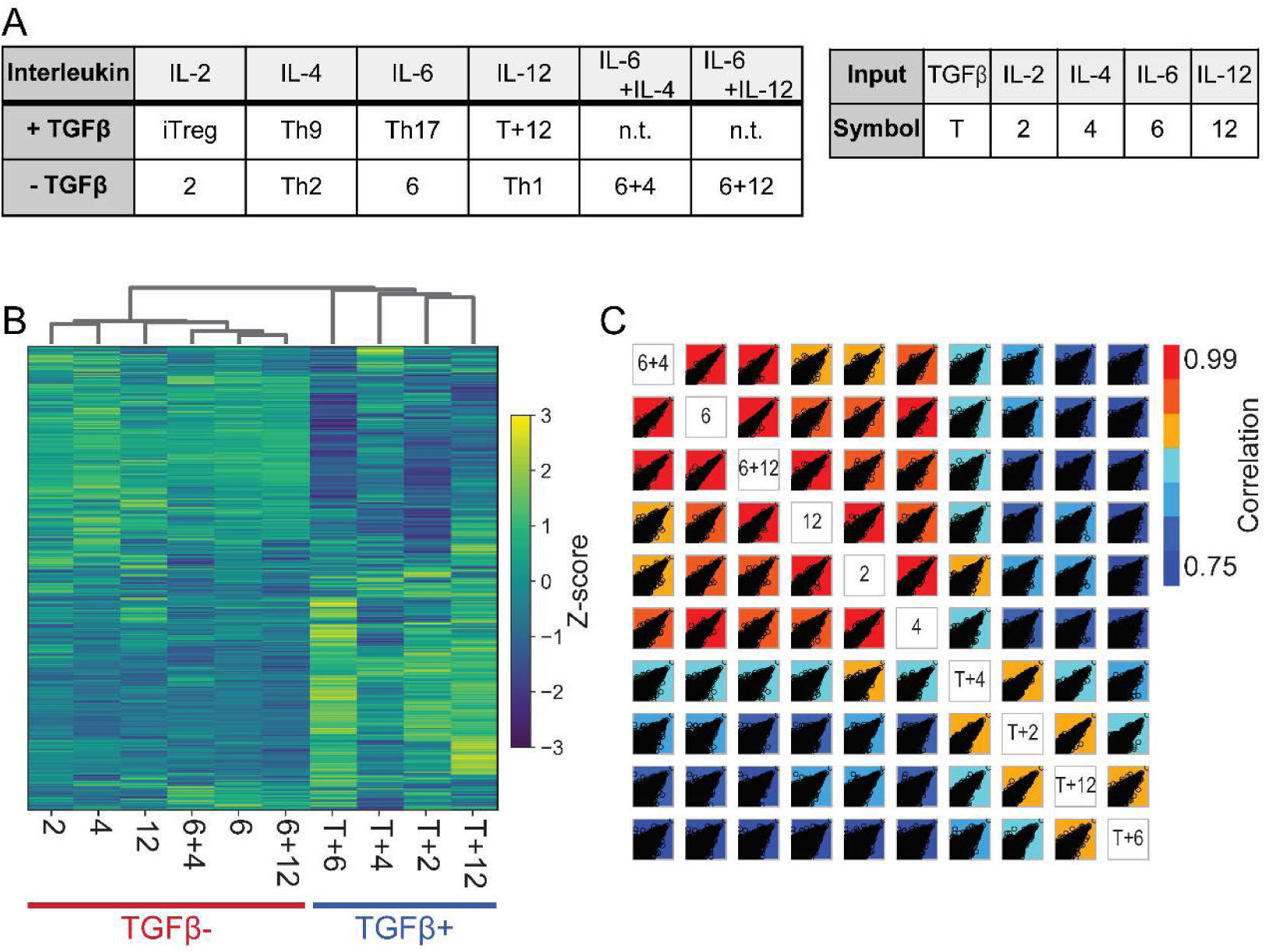
CD4^+^ T cells display distinct mRNA profiles following culture in the presence or absence of TGFβ in combination with other cytokines. Naïve murine CD4^+^ T cells were cultured with stimulation using plate-bound anti-CD3 and anti-CD28, together with different cytokines. After 96h, mRNA was extracted and used for genome-wide gene expression analysis using RNA-seq. (A) Left: The cytokine combinations used in this study. Rows and columns indicate the applied cytokine combination, and values indicate the T-cell state that is induced by the specific combination. (n.t: not-tested). Right: Samples name abbreviations. (B) Heatmap showing the expression levels of 11,308 genes, clustered by standardized gene expression levels (Z-score). Clustering of samples according to the correlation between their gene expression profiles is shown above the heatmap (C) Correlation matrix indicating the level of similarity in gene expression between all pairs of samples. The color of each box represents the correlation coefficient (color bar). Genes are represented by black dots, showing the levels of expression of all genes in the two conditions. Results are averaged over n = 3 repeats. See also Figure S1.

After removing lowly-expressed genes, we clustered the remaining 11,308 genes according to their level of expression (Fig. 1B). We found that samples cluster according to the presence or absence of TGFβ as an input cytokine (Fig. 1B, Fig. S1A). In the absence of TGFβ as an input, the samples containing IL-6 clustered together and differed from samples without IL-6 (Fig. 1B, Fig. S1A). This hierarchy in the overall influence of cytokines is in accordance with our previous findings (17). Samples in which TGFβ was present as an input were highly correlated between themselves and showed a lower correlation with samples that did not contain TGFβ (Fig. 1C). The sample of TGFβ+IL-4 lies in between the two groups, suggesting an intermediate phenotype of these cells (Fig. 1C). Differential expression analysis identified 1,177 genes that were significantly altered by TGFβ (either up-or down-regulation; fold-change>2, adjusted p-value<0.05). As a comparison, IL-6 significantly influences only 116 genes, an order of magnitude fewer.

### TGFβ affects the expression of genes associated with CD4^+^ T-cell differentiation, metabolism, cell death, and localization

In order to validate the known effects of cytokine combinations, we examined the expression of genes that characterize the main CD4^+^ T-cell subsets (Fig. 2A). As expected, lineage-specific transcription factors (TFs) such as *Tbx21* (encoding T-bet, Th1), *Gata3* (Th2), *Rorc* (Th17), and *Foxp3* (Treg) displayed high expression levels in samples exposed to cytokines known to induce differentiation into the corresponding lineage (Fig. 2A, Fig. S1B). A similar trend was also found for lineage-associated cytokines and their receptors. Additionally, these phenotype-related genes were expressed at intermediate levels in other cytokine combinations, in agreement with our previous results showing a continuum of intermediate states (17, 18). For instance, *Foxp3*, which is considered a Treg lineage-specific TF, had the highest expression in the Treg-inducing condition, TGFβ+ IL-2. However, *Foxp3* was also expressed in other samples at intermediate levels. Significantly, all samples cultured with TGFβ showed higher expression of Foxp3 compared with samples where TGFβ was not added (Fig. 2A, Fig. S1B).

**Figure 2.**
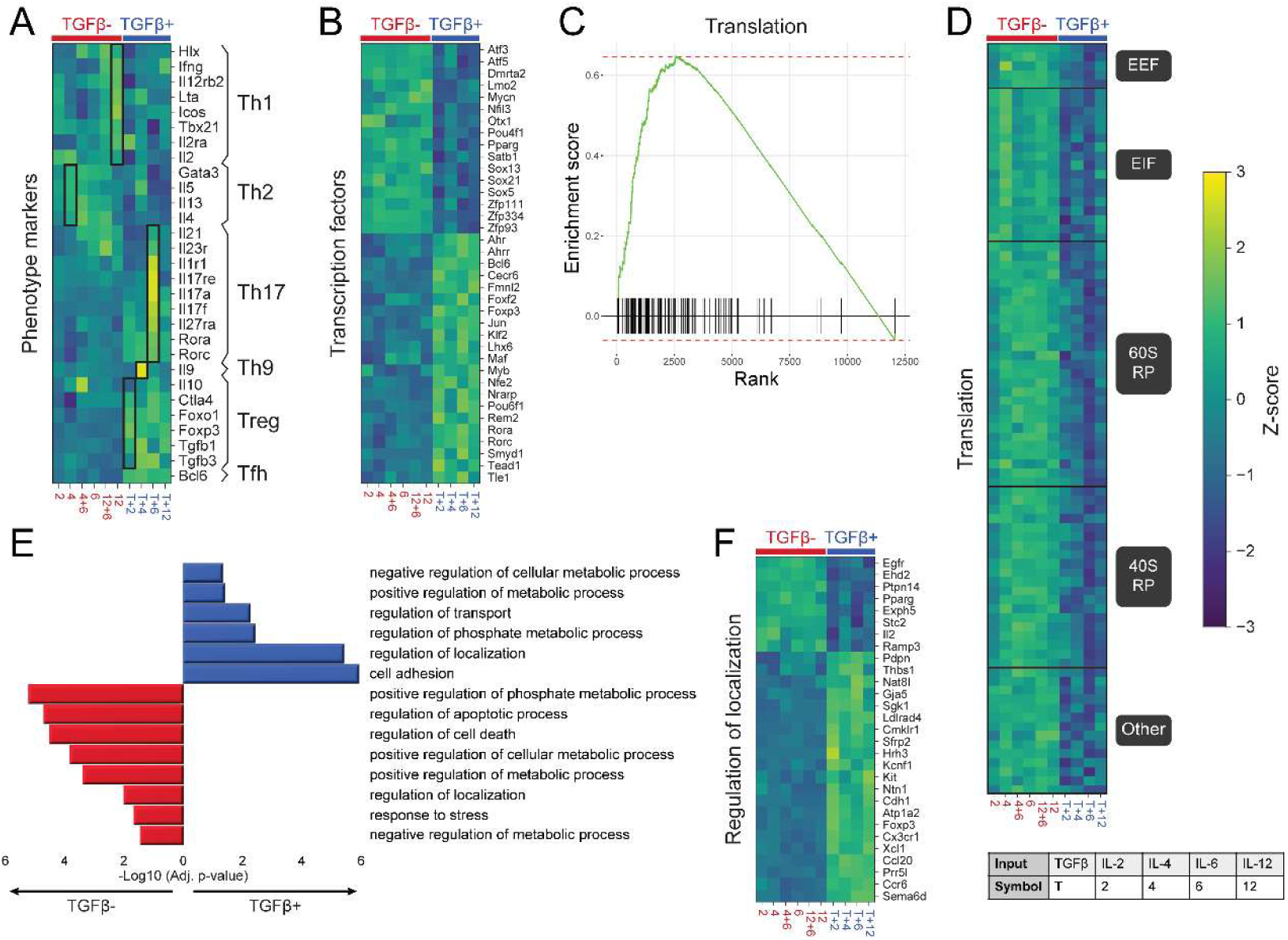
TGFβ, in combination with other cytokines, induces changes in genes associated with CD4^+^ T-cell differentiation, metabolism, localization, and cell death. (A) Normalized expression levels of known lineage-specific genes, characterizing different CD4^+^ T-cell phenotypes. Black boxes mark conditions that drive polarized Th fates and the corresponding characterizing genes. (B) Normalized expression levels of TFs that were differentially expressed between the TGFβ^+^ and TGFβ^-^ groups (Fold Change>2; SD≥0.8; *P*-value≤0.05, see Methods). (C and D) Gene-set enrichment analysis (GSEA) showing the overrepresentation of genes involved in protein translation in the TGFβ^-^ group. (C) Enrichment score for the “Translation” gene set, showing an overrepresentation of translation-related genes in the TGFβ^-^ group. (D) Expression level heatmap of the GSEA leading-edge subset of the “Translation” gene set. (E and F) Gene ontology (GO) enrichment analysis of differentially expressed genes between samples with IL-2 or IL-12 as input cytokines and the samples with IL2+TGFβ or IL12+TGFβ. (E) List of selected biological process gene ontology (GO) terms and their significance score (FDR-adjusted p-value; blue: enriched in the TGFβ^+^ group; red: enriched in the TGFβ^-^ group). (F) Normalized expression levels of selected genes associated with regulation of localization (Fold Change>4; SD≥0.8; *P*-value≤0.05). Cytokine combinations are marked using one-symbol labels, as specified in the table at the bottom. Results are averaged over n = 3 repeats. See also Figure S1.

Next, we examined which TFs are differentially expressed between samples that received TGFβ as an input (TGFβ^+^ group) and samples that did not receive TGFβ as an input cytokine (TGFβ^-^ group) (Fig. 2B). Several identified TFs had previously been linked to CD4^+^ T-cell differentiation. For example, aryl hydrocarbon receptor (AHR) and aryl hydrocarbon receptor repressor (AHRR), which showed higher expression levels in the TGFβ^+^ group, are known to be involved in central processes in Treg and Th17 cells (19–21). KLF2 (high in TGFβ^+^) was found to control the differentiation of induced Tregs and Treg migration (22, 23). NFIL3 (high in TGFβ^-^) was shown to be a regulator of key TFs and cytokines in a number of CD4^+^ T-cell subsets (24, 25), including Th1 (25), Th2 (26), Th17 (27), and Treg (28). We note that in some cases, the changes we observe seem contradictory to previous observations. For instance, PPARg, which we found to be high in the TGFβ^-^ group, was shown to be a driver of Treg function (29–31) and also to restrict Th17 differentiation through repression of RORγt (32). Overall, TGFβ robustly affects a global set of transcription factors associated with CD4^+^ T cells.

Further examination revealed that *Itgae*, the gene encoding the tissue-resident memory T cells (T_RM_) marker CD103, displayed much higher expression in cells primed under all four cytokine mixtures that include TGFβ (Fig. S1C). Moreover, we found that many genes that are involved in the differentiation and maintenance of T_RM_ cells (33) were also differentially expressed in our data and showed higher levels in TGFβ^+^ conditions compared with TGFβ^-^ conditions (Fig. S1D). These include *Cxcr6*, *Cx3cr1*, *Cd101*, *Itgb2*, and *Klf2*, among other genes. Furthermore, we found that activation of CD4^+^ T cells in the presence of cytokine combinations that include TGFβ resulted in the expression of *Cd8a*, which encodes the hallmark surface marker of cytotoxic T cells (Fig. S1E). TGFβ^+^ conditions also lead consistently to the elevation of *Ctsw* (Fig. S1E), which encodes the protein Cathepsin W that is believed to be restricted to cytotoxic T cells and NK cells and is thought to have a role in T-cell cytolytic activity (34, 35). Collectively, TGFβ affects diverse aspects of the resulting phenotype of the cells, beyond controlling the CD4^+^ T helper subsets.

To further study the effect of TGFβ on gene expression, we examined the association of differentially expressed genes to biological processes. Gene-set enrichment analysis (Fig. 2C, D) revealed higher levels of genes related to translation in cells in the TGFβ^-^ group compared with the TGFβ^+^ group. Specifically, most ribosomal proteins from both 40S and 60S subunits were expressed at lower levels when TGFβ was present, as well as many translation initiation and elongation factors (Fig. 2D, Fig. S1F). Gene ontology (GO) analysis revealed enrichment also for genes related to apoptosis and cell death in the TGFβ^-^ group (Fig. 2E, Fig. S1G). Intriguingly, genes related to localization (Fig. 2F) and metabolic processes (Fig. S1H) were enriched in cells from both TGFβ^-^ and TGFβ^+^ groups; however, the specific genes associated with these functionalities differed between the two groups. Taken together, our findings suggest high translational activity in T cells activated by αCD3+ αCD28 in the presence of stimulatory cytokines.

The cytokine combinations that we studied are typically investigated with respect to their diverse influence on CD4^+^ T-cell differentiation. Our analysis reveals that TGFβ, when present with other cytokines, modulates the expression of a large set of genes that are involved in many processes of T-cell physiology, including translational activity, cellular death, localization, and metabolism. Notably, the gene expression pattern clusters cells based on their exposure to TGFβ and not based on their regulatory and effector phenotypes.

### TGFβ exposure gives rise to smaller CD4^+^ T cells in vitro and in vivo

Following activation, naïve CD4^+^ T cells dramatically increase their volume, a process that requires metabolic adaptation and enhanced translation of proteins (36–39). The size of lymphocytes is largely determined by the rate of protein synthesis, whose increase requires ribosome biogenesis (40). Our gene expression analysis revealed that, during T-cell activation, TGFβ exposure modulates the expression of genes associated with translation and other metabolic processes. This suggests a potential effect of TGFβ on the size of cells.

When examining the forward-scatter (FSC) and side-scatter (SSC) values of cells, parameters that are routinely measured by flow cytometry and relate to cell size and morphology, we found them to be affected by the cytokine milieu (Fig. 3A). Analyzing our previously published dataset of 64 cytokine combinations (17) we find that conditions that include TGFβ induced smaller FSC and SSC values compared with conditions that lack TGFβ (Fig. 3B). Lower FSC values suggest that cells growing in the presence of TGFβ are of a reduced size. We have previously developed a multi-parameter linear regression model capable of predicting CD4^+^ T-cell differentiation markers from the cytokine milieu (17). Here, we extended the model predictions beyond the biochemical outcomes and found that the same model can capture the dependence of physical related measurements such as FSC and SSC values on the input cytokine combination (Fig. 3B inset and Fig. S2A, B).

**Figure 3.**
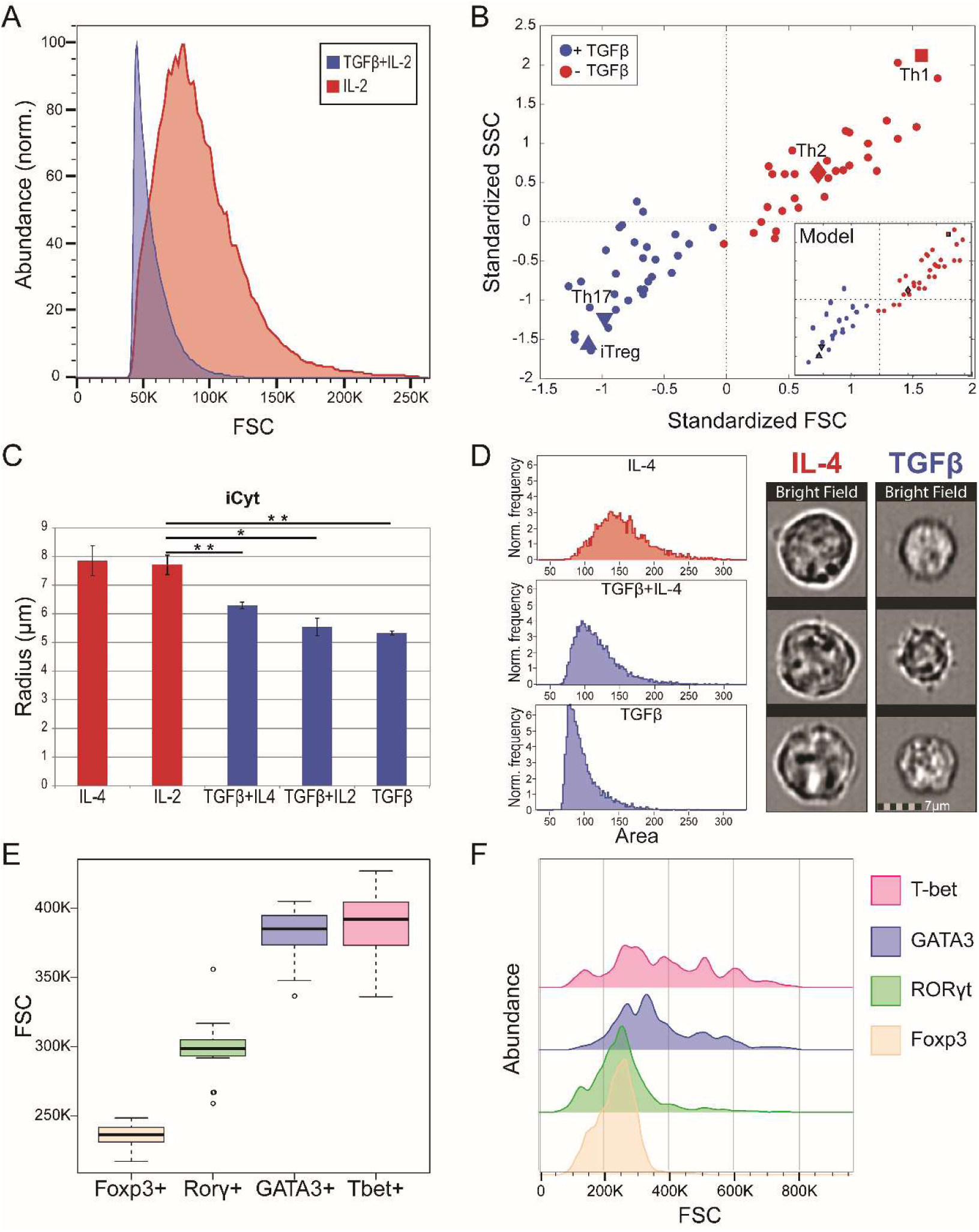
TGFβ affects the size of in vitro and in vivo activated CD4^+^ T-cells. The size of CD4^+^ T cells was examined both in vitro (A-D) and in vivo (E-F) as following: (A-D) Naïve CD4^+^ T cells were isolated and cultured under different cytokine combinations, with and without TGFβ. (A) FSC measurements of two representative conditions: with TGFβ (TGFβ+ IL-2, blue) and without TGFβ (IL-2, red). (B) Mean FSC and SSC values were measured under 64 different cytokine combinations and standardized to have zero mean and unit standard deviation. Samples are colored according to the presence (blue) or absence (red) of TGFβ in the culture medium. Conditions that drive polarized Th lineages are marked. The mean values of three independent experiments were used in the analysis. The predicted response of the cells to the 64 cytokine combinations, as described by a mathematical model that we have previously developed (17) is shown (inset). (C) iCyt measurements of cell radius under different cytokine conditions (n=3). Bars are colored according to the presence (blue) and absence (red) of TGFβ as an input cytokine. (D) Area of activated cells (CD4^+^CD44^hi^), as measured using imaging flow cytometry (ImageStreamX). Left: representative cell area distributions for the specified conditions. Right: representative images of cells (bright field) cultured with IL-4 or with TGFβ. (E-F) Size of CD4^+^ T cells in vivo. Mice were vaccinated with influenza virus+adjuvant (see Methods). Spleen CD4^+^ T cells were collected 48h following boost and analyzed by flow cytometry to define their differentiation state by transcription factor expression: T-bet (Th1), GATA3 (Th2), RORγt (Th17), and Foxp3 (Treg). FSC levels of the gated populations were measured using flow cytometry as an indicator for cell size. (E) FSC measurements of in vivo activated CD4^+^ T cells. Data is pooled from two independent experiments, n= 7 mice per experiment. (F) Representative flow cytometry plot showing the FSC expression on CD4^+^ T cells expressing Foxp3^+^ (orange), RORγt^+^ (green), GATA3^+^ (blue), or T-bet^+^ (pink). See also Figure S2.

We supplemented this flow cytometry analysis by including additional experimental approaches to measure the size of cells with and without TGFβ. To directly measure cell size, we first used electronic measurements of cell volume (EV) based on the Coulter method (see Methods). Our analysis showed that cells grown in the presence of cytokine combinations that contain TGFβ displayed a significantly smaller volume (Fig. 3C, Fig. S2A). As EV linearly correlates with FSC (Fig. S2B), we use FSC values, which are more readily measured, as a proxy for cell size.

To focus on size difference that does not arise from the activation state of the cells, we further examined the effect of TGFβ on the size of activated CD4^+^CD44^+^ T cells. Using Imaging Flow Cytometry, we found that when cultured with cytokine mixes that include TGFβ, activated cells were consistently smaller and had a narrower size distribution compared to cells cultured in the absence of TGFβ (Fig. 3D, S2C).

Furthermore, cells activated in the presence of TGFβ alone were even smaller and less granular (Fig. 3C, D).

To investigate differential CD4^+^ T-cell sizes in vivo, we immunized mice with influenza virus. Mice were boosted on day 14, and 2 days later cells were isolated from spleens. Isolated cells were stained to identify the CD4^+^ T cells population and determine their activation state and expression of lineage-specific TFs. CD4^+^CD44^+^ T cells expressing T-bet (Th1) or GATA3 (Th2) displayed significantly higher FSC values compared to RORγt^+^ (Th17) or Foxp3^+^ cells (Tregs) (Fig. 3E, F). Similar results were obtained for cells isolated from draining lymph nodes (Fig. S2D). Collectively, we establish that activated CD4^+^ T-cell subsets that require TGFβ for their differentiation display significantly smaller sizes both in vitro and in vivo.

### TGFβ affects the size of activated proliferating CD4^+^ T cells

We next examined whether the effect of TGFβ on the size of CD4^+^ T cells depends on their proliferation. Naïve CD4^+^ T cells were stained with a cell-proliferation dye (CFSE) and cultured with activation under different cytokine conditions, with or without TGFβ. Cell proliferation and survival were examined at intervals of 24h, for the duration of 4 days, using flow cytometry. We observed that under all conditions tested, some cells did not proliferate even after 3-4 days of culture, in agreement with previous reports (41, 42). These non-dividing cells were of similar small size, regardless of the cytokine combination that they were exposed to (Fig. 4A). Conversely, only proliferating cells modulated their size in response to cytokine mixtures, and in particular, were smaller in the presence of TGFβ than in its absence. We then examined the size of proliferating cells in different generations following 72h and 96h of culture with IL2 or with IL2+TGFβ. As before, evident differences in size are observed between cells cultured in the presence or absence of TGFβ as an input cytokine. However, the size of the cells was similar in different generations in both conditions and both time points (Fig. 4B). Therefore, the presence of TGFβ affects the size of dividing cells, regardless of their generation number, and does not affect the size of non-proliferating cells.

**Figure 4.**
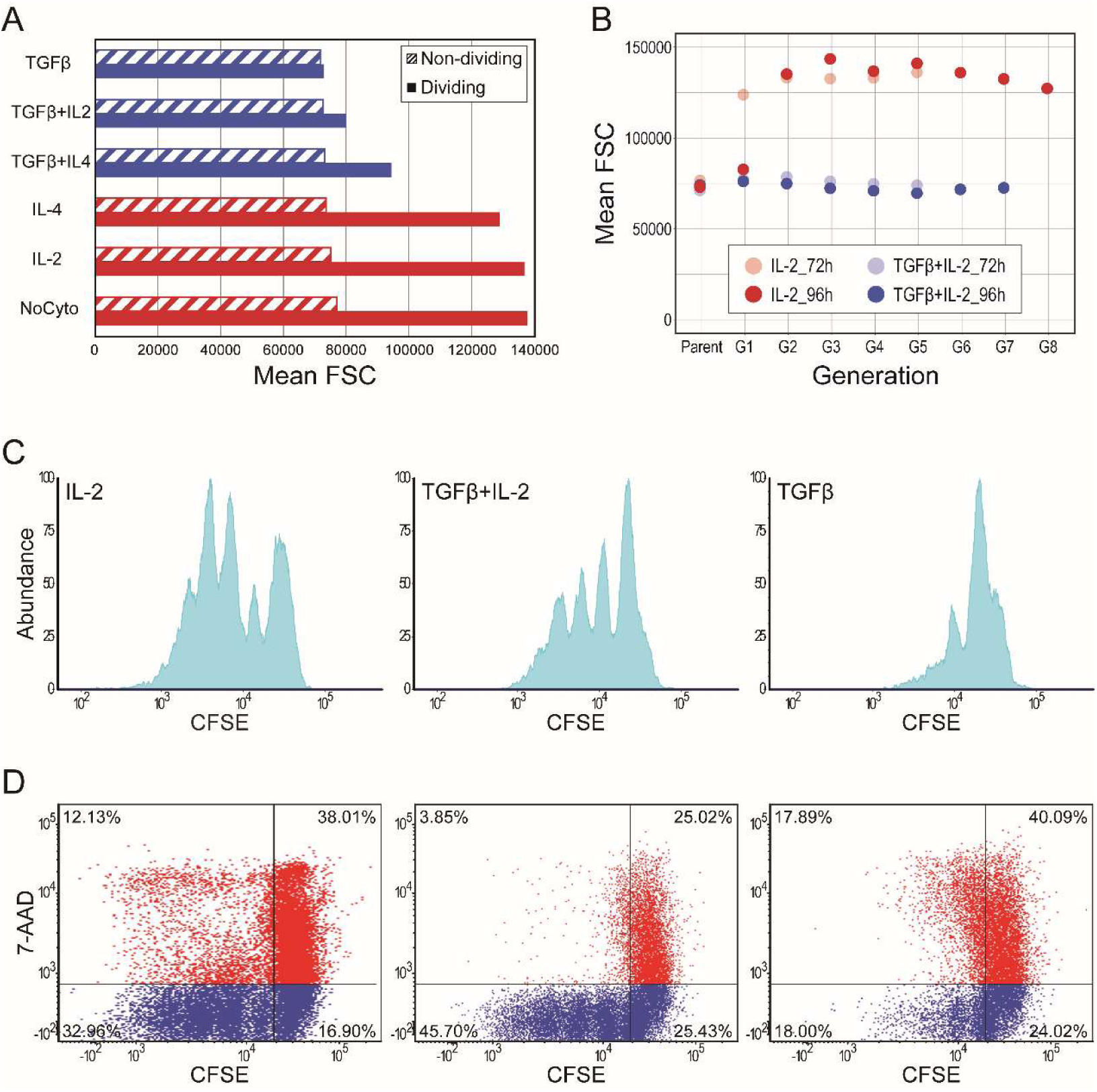
TGFβ influences the proliferation and survival of CD4^+^ T cells. Naïve CD4^+^ T cells were isolated and cultured under different cytokine combinations, as indicated. Proliferation and death of the cells were examined in intervals of 24h. (A) Mean FSC of dividing and non-dividing cells under different cytokine conditions. (B) Mean FSC of cells in different generations, as determined by gating on CFSE labeling. (C) Proliferation profiles of three representative conditions: IL-2 (left), TGFβ+ IL-2 (middle), and TGFβ (right) following 96h of culture. (D) Scatter plots showing cell death (by 7-AAD staining) and proliferation (by CFSE) following 72h of culture under three representative conditions: IL-2 (left), TGFβ+IL-2 (middle), and TGFβ (right). See also Figure S3.

To further examine the proliferation and survival profiles of the cells, we compared three conditions: culture with TGFβ, culture with IL2, and culture with both cytokines (Fig. 4C, D). In the presence of TGFβ alone, the majority of cells did not divide, and proliferating cells underwent a limited number of divisions (Fig. 4C, D right panel, Fig. S3A), in accordance with previous observations (43–45). When exposed to IL2 alone, cells displayed a high proliferation rate, combined with a profound death of both proliferating and non-proliferating cells (Fig. 4C, D left panel). When both TGFβ and IL2 are present, substantial proliferation is observed (Fig. 4C, D middle panel). The maximal number of divisions was similar to that obtained with IL2 alone, but the proliferation profile was distinct, with fewer cells reaching maximal proliferation. In the presence of TGFβ, dividing cells were mostly viable, in contrast with the higher death rates observed with IL2 alone. These observations repeated themselves in the combinations of TGFβ + IL4 (Fig. S3B, C). Together, we find that in the presence of additional cytokine signals, TGFβ does not prohibit CD4^+^ T-cell proliferation but significantly increases the survival of dividing cells.

### A dense network of interacting transcription factors links TGFβ signaling with cell growth

To identify candidate genes that mediate the effect of TGFβ on cell size, we generated a regulatory network connecting TGFβ and TCR activation to cell growth. We started by analyzing all TFs that were differentially expressed in response to TGFβ according to our RNA-seq data (Fig. 2B). To study the effect of TGFβ beyond guiding cell differentiation, we removed TFs that have a known role in T-cell fate determination. Using String (46), we extracted the network of interactions between the remaining TFs, and further extended the network to include additional connected genes while discarding all disconnected nodes. Finally, using this list and based on an extensive literature search (Table S4), we generated a curated network of established interactions (Fig. 5). Overall, we obtained a dense network that includes signaling molecules and TFs, which are influenced by TGFβ in the context of CD4^+^ T-cell activation, and contribute to the control of key cellular processes including division, survival, and growth.

**Figure 5.**
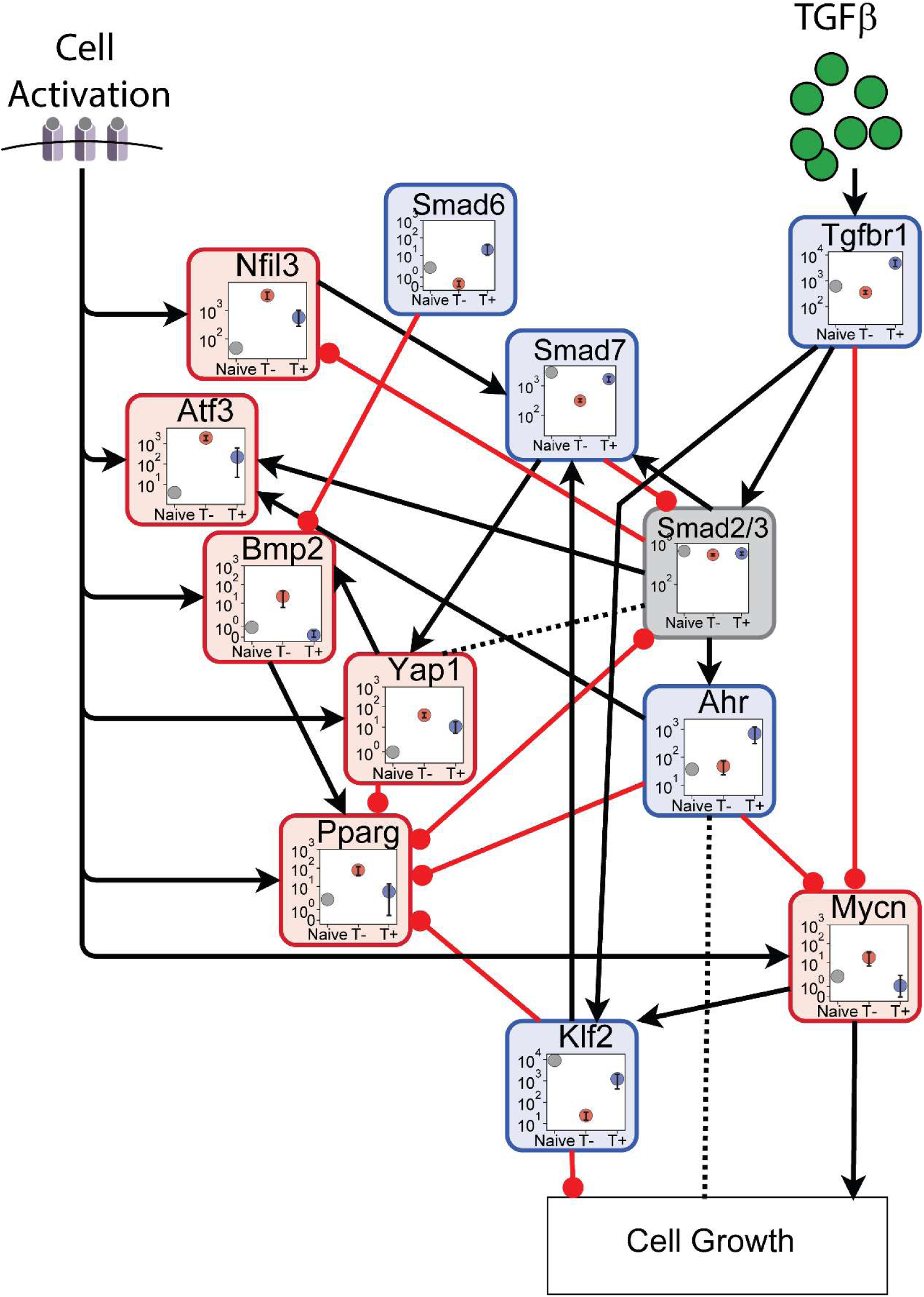
Gene interaction network controlling cell growth in response to TGFβ. A curated network of known interactions between TFs that are differentially expressed in CD4^+^ T cells in the presence of TGFβ, and that are potentially involved in the regulation of cell size. Each node displays normalized expression levels of the TF in naïve T cells (“Naïve”, gray), or in T cells that were activated in the absence (“T-”, red) or presence (“T+”, blue) of TGFβ. The color of each node represents whether gene expression levels are higher (blue), lower (red), or similar (gray) in the presence of TGFβ. Solid black arrows represent induction or activation, while solid red arrows represent inhibition or repression. Dashed links represent more complex crosstalk between the connected nodes. In the SMAD 2/3 node, the expression of SMAD2 is shown. Network links are supported by references shown in Table S4.

The network of genes we identified suggests possible candidate genes that mediate TGFβ signals to control cell growth. Specifically, both *Klf2* and *Mycn* appear to serve as key nodes regulating cell growth, and are controlled by TGFβ both directly and indirectly through a dense interconnected genetic network (Fig. 5). TGFβ increases the levels of *Klf2*, which in turn inhibits cell growth, while the levels of *Mycn*, a positive regulator of cell growth, are decreased in the presence of TGFβ. Thus, our curated literature analysis has identified *Klf2* and *Mycn* as possible candidate genes that mediate the TGFβ control of cell size.

### TGFβ affects the expression levels and localization of N-Myc

To test the hypothesis arising from the network analysis, we used qPCR to examine the expression levels of *Klf2* and *Mycn* under different cytokine conditions with and without TGFβ (Fig. 6A). Similar to the RNA-seq data, the expression of *Mycn* was reduced in the presence of TGFβ, while the expression of *Klf2* was enhanced (Fig. 6A). In addition, we compared the expression of *Mycn* to its paralog *Myc*, which does not appear in our network but has been associated with T-cell growth and proliferation (40, 47, 48). Although both *Mycn* and *Myc* are generally known to be inhibited by TGFβ (49–52), we found that during CD4^+^ T-cell activation, these TFs show different expression patterns (Fig. 6A). While *Mycn* expression depended on TGFβ, *Myc* was expressed at similar levels regardless of the presence of TGFβ in the cytokine mixture.

**Figure 6.**
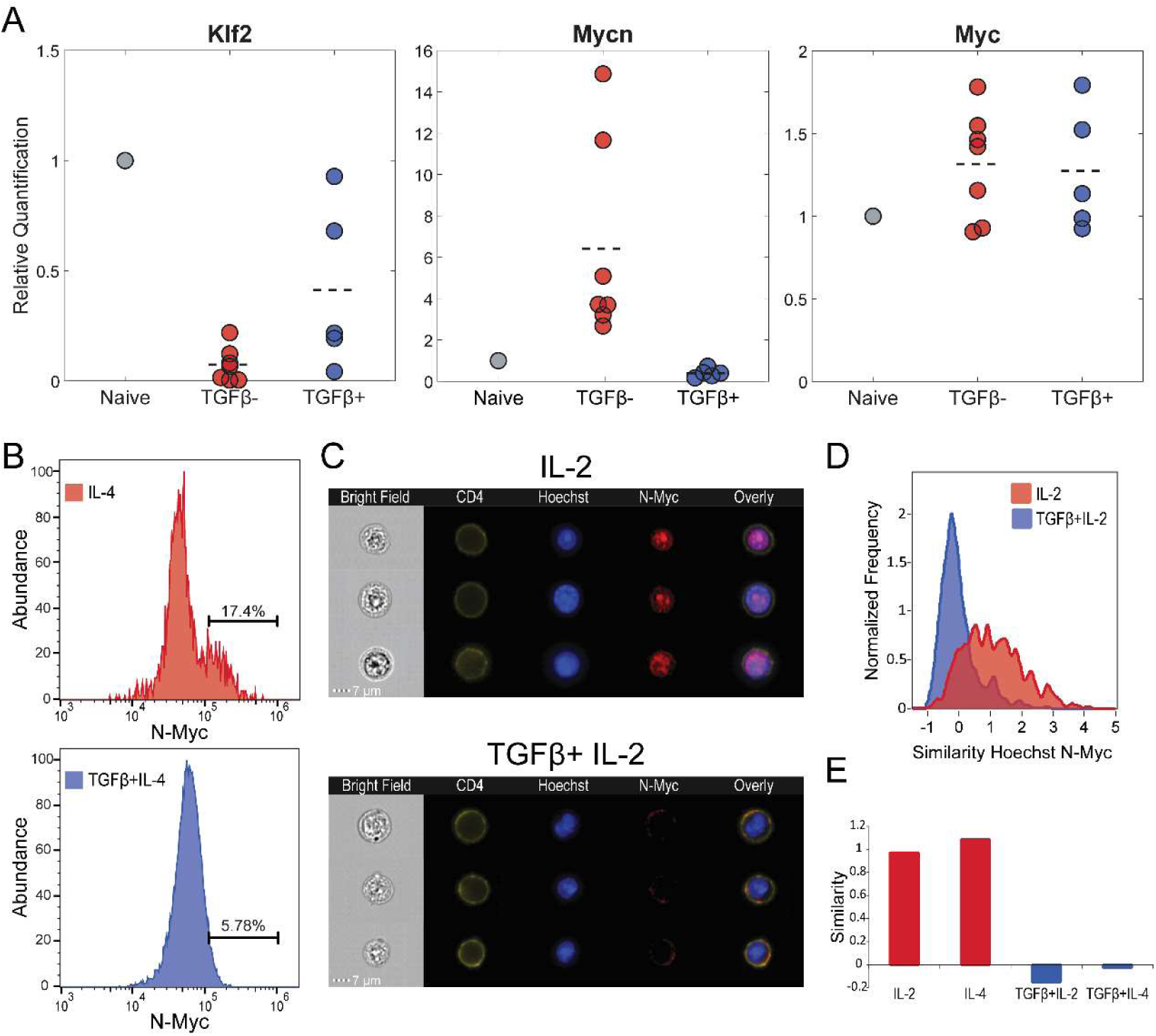
Gene and protein expression and localization in the presence and absence of TGFβ. (A) Expression levels of: *Klf2* (left), *Mycn* (middle), and *Myc* (right) in CD4^+^ T cells cultured under different cytokine conditions, with (blue) and without (red) TGFβ. Naïve cells sample (grey) is also shown for comparison. Expression levels were examined using qPCR. Data represent means (n=3). (B-E) Protein expression levels and localization of N-Myc were examined following 96h of culture, in the presence or absence of TGFβ, using imaging flow cytometry. Cells were stained for: N-Myc, CD4 (membrane), and Hoechst (nucleus). (B) Histograms showing the expression levels of N-Myc in cells cultured with IL4 (red) and with TGFβ+IL4 (blue). (C) Images of six representative cells, three cultured with IL-2 and three cultured with TGFβ+IL-2. Shown are five images for each cell: bright field (sample illustration, left), CD4 staining (cell membrane, 2nd left), Hoechst staining (nucleus, center), intracellular N-Myc staining (2nd right), and their composition (right). (D) Histograms showing the level of localization of N-Myc to the nucleus, as measured by the overlap between Hoechst and N-Myc staining, under two culture conditions: IL-2 (red) and TGFβ+IL-2 (blue). (E) Mean level of similarity between Hoechst (nucleus) and N-Myc, under four culture conditions.

Further studying the possible role of *Mycn* in the TGFβ-dependent control over cell growth, we tested at the protein level how TGFβ signals affect the response of N-Myc to TCR activation. We used imaging flow cytometry to examine N-Myc protein expression and subcellular localization in cells cultured in the presence or absence of TGFβ. As observed for mRNA, we found that the expression of the N-Myc protein was higher in the absence of TGFβ (Fig. 6B). Moreover, we found that TGFβ also affected the nuclear localization of N-Myc, thus critically modifying its functionality as a transcription factor. While N-Myc was found inside the nucleus in cells activated in the presence of IL2 alone, it was excluded from the nucleus in the presence of TGFβ+IL2 (Fig. 6C).

To obtain a quantitative measure for the localization of N-Myc, we stained the cultured CD4^+^ cells with a membrane marker (CD4) and a DNA stain (Hoechst) (Fig. 6C), and calculated the level of similarity between the expression of N-Myc to that of these two markers. We found high similarity between N-Myc and the nuclear stain in CD4^+^ T cells activated in the presence of IL2, and significantly lower similarity in the presence of TGFβ+IL2 (Fig. 6D). Similar results were obtained when IL4 was compared with IL4+TGFβ (Fig. 6E). Collectively, we established that TGFβ modulates N-Myc activity by both decreasing its level and excluding it from the nucleus, thus preventing its interaction with gene regulatory elements.

## Discussion

TGFβ is a pleiotropic cytokine that affects many cellular functions in a large number of cell types. However, while TGFβ is known to affect diverse aspects of cellular state in various biological contexts, much is still unknown about its role in CD4^+^ T cells, beyond guiding their differentiation. Here, we distilled the effect of TGFβ from that of other signals by analyzing responses that consistently depend on TGFβ, regardless of the other cytokines in the mix. Using this approach, we comprehensively investigated the effects of TGFβ on multiple aspects of CD4^+^ T cells state.

Gene expression profiling revealed many genes that were consistently upregulated or repressed by TGFβ. TGFβ affected not only the differentiation state of the cells (i.e., their immune phenotype) but also other characteristics, such as cell size, proliferation, and death. Our RNA-seq analysis further revealed an enrichment for localization and metabolic processes in the presence of TGFβ. Intriguingly, such enrichment is observed in both upregulated and downregulated genes, suggesting a TGFβ-dependent shift in the immunometabolic state of the cells. One can assume that these unique metabolic properties are tailored to support the phenotype and immune functions of the different T-cell fates, characteristics which are highly influenced by the cytokine environment. New approaches, such as the characterization of the metabolic state of cells at the single-cell level (53) and studying T-cell metabolism under in vivo settings (54), would provide more in-depth insights into the effect of TGFβ on the metabolic profile of the cells. Such understanding could open up new and promising therapeutic approaches by enabling the modulation of T-cell phenotypes through manipulations of cellular metabolism.

When examining cell morphology, we found that cells cultured in the presence of TGFβ exhibited a reduced size, both in vitro and in vivo. TGFβ affected the size of activated and proliferating cells but not that of non-dividing cells. Accordingly, genes associated with translation were expressed at lower levels when TGFβ was present in the cytokine mixture. As cell size and growth are generally associated with metabolic adaptations, this raises the question of whether the metabolic changes induced by TGFβ are required to control cell size.

In addition, we found several genes important for the generation and maintenance of tissue-resident memory T cells (T_RM_) to be elevated in TGFβ^+^ conditions. Therefore, priming of naïve T cells in the presence of TGFβ together with stimulatory cytokines could promote the formation of a cell population that is equipped for tissue residency. The smaller size of these cells may facilitate traffic within the dense tissue environment, and the expression of genes that affect localization and adhesion can shape their location and maintenance within tissues. As cells migrate to and reside in distinct tissues, they are being exposed to different nutrients and metabolites and must adapt to the tissue ecosystem, with its available resources. A tempting hypothesis might be that in parallel to the elevation of characteristic T_RM_ genes, TGFβ also introduces metabolic changes in cells, equipping them to utilize available nutrients and metabolites in tissues they could potentially inhabit.

By combining an extensive literature search with our mRNA measurements, we were able to highlight a network of genes whose transcription is modulated by TGFβ and that have been previously associated with cell growth. The resulting network highlighted *Mycn* as a focal point between TGFβ-dependent signals and the control of cell growth. The Myc gene family of transcription factors was shown to have a role in regulating cell growth (48, 55, 56). Both c-Myc and N-Myc, two members in the Myc gene family, have putative target genes in cell growth control (48, 57). Moreover, it was previously demonstrated that the expression of c-Myc rapidly increases within several hours in response to IL-2 (58) or TCR ligation (59). And finally, TGFβ is known to inhibit both c-Myc (49, 50) and N-Myc (51, 52, 60, 61). Indeed, we found *Mycn* expression, as well as N-Myc protein levels, to differ between cells cultured in the presence and absence of TGFβ. Consistently, by analyzing the data collected in a previous comprehensive study of TFs expression in different populations of CD4^+^ T-cell phenotypes (62), *Mycn* displayed lower expression levels in cells cultured under TGFβ^+^ conditions. Furthermore, we found that N-Myc was excluded from the nucleus in the presence of TGFβ. These results indicate that TGFβ prevents N-Myc from acting as a functional transcription factor. Taken together, our observations suggest N-Myc as a possible mediator that is regulated by TGFβ both at the expression level and at its intracellular localization, and could be a core element mediating the regulation of TGFβ on T-cell size.

Overall, we have found that TGFβ plays a significant role in many cellular processes, including differentiation, survival, proliferation, and cell size. Since TGFβ is widely expressed in many tissues and physiological contexts, it will be important in future work to carefully dissect the mechanistic relationships between these effects. Our findings establish an in vitro culture system to study cell size control. Exposing cells to systematic combinations of two or more stimuli, including growth factors, membrane receptor/ligand or a combination of the two, offers a powerful approach to understanding how cells respond in the rich, diverse, and complex microenvironments that exist in vivo.

## Acknowledgments

We are grateful to Prof. B. Chain for his critical reading of the manuscript and helpful discussion. N.F. was supported by the Applebaum Family Foundation. Y.E.A is supported by the Israel Science Foundation (grant 1105/20) and is the incumbent Sygnet Career Development Chair for Bioinformatics.

## Data availability

Bulk RNA sequencing data have been deposited at GEO: GSE264451 and are publicly available.

