## Supplemental Data 1 for "A dominant role of TGFβ in regulating T-cell size and physiology"

**Experimental Procedures**

**Mice.** C57BL/6 female mice were purchased from Harlan Laboratories. Mice were housed under specific pathogen-free conditions at the animal facility of the Weizmann Institute and used at 6–8 wk of age. For in vivo polarization experiments, C57BL/6 male mice at the ages of 8-10 wk were used. All animal experiments were performed under protocols approved by the Animal Care and Use Committee of the Weizmann Institute.

**Cell Culture.** Naïve $CD4^+$ T cells were isolated from spleens of mice by magnetic microbeads separation (CD4+CD62L+ MACS T cell isolation kitII, Miltenyi Biotec) and MACS separation columns, according to the manufacturer's instructions. Cells were cultured at a concentration of $1x10^6$ cells/ml, in the presence of plate-bound anti-CD3 (1µg/ml) and anti-CD28 (3µg/ml) for stimulation. Cells were cultured in complete RPMI-1640 medium with phenol red, supplemented with: 10% (vol/vol) fetal calf serum (FCS), 2mM L-Glutamine, 1% Penicillin-Streptomycin, 1mM Sodium Pyruvate, 1% Kanamycin, 1% non-essential amino acids, and 50µM β-mercaptoethanol. Cells were supplemented with different combinations of cytokines, used in the following concentrations: 10ng/ml rhTGFβ1 (R&D #240-B-002), 5ng/ml rmIL-2 (R&D #402-ML-020), 20ng/ml rmIL-4 (R&D #404-ML-010), 20ng/ml rmIL-6 (R&D #406-ML-005) and 10ng/ml rmIl12 (R&D #419-ML-010). Cells were cultured for 96 hours and were measured as detailed below.

**Cell Proliferation Assay.** Cells were collected as specified before. Before culturing, naïve cells were labeled with 2µM carboxyfluorescein diacetate succinimidyl ester (CFSE; Invitrogen) for 10 min at 37°C. After incubation, cells were washed with PBS, resuspended in complete RPMI media, and cultured as described above. Cell proliferation was examined in intervals of 24h. Prior to flow cytometry analysis, 7-AAD (Biolegend) was added to the cells to distinguish live cells from the dead. Cell proliferation was assessed by CFSE dilution using flow cytometry (LSRII, BD) and data were further analyzed with FlowJo software (TreeStar) and FCS Express (De Novo Software).

**RNA Preparation, Library Construction, and RNA Sequencing.** Following 96h of culture, cells went through ficoll, and total RNA was isolated using NucleoSpin RNA kit (MACHEREY-NAGEL), according to manufacturer's instructions. Libraries were prepared using the INCPM-mRNA-seq. Briefly, the polyA fraction (mRNA) was purified from 300-400 ng of total input RNA followed by fragmentation and the generation of double-stranded cDNA. After Agencourt Ampure XP beads cleanup (Beckman Coulter), end repair, A base addition, adapter ligation and PCR amplification steps were performed. Libraries were quantified by Qubit (Thermo fisher scientific) and TapeStation (Agilent). Sequencing was done on a Nextseq 75 cycles high output kit, allocating 15M reads per sample (Illumina; single read sequencing).

**Quantitative RT-PCR.** Following 96h of culture, cells went through ficoll, and total RNA was isolated using RNeasy micro kit (Qiagen) for all samples besides the naïve cells, for which the RNeasy mini Kit (Qiagen) was used, according to manufacturer's instructions. Isolated RNA was used to synthesize cDNA using M-MLV reverse transcriptase (Promega) for the detection of Klf2, and using Lunascript RT Supermix Kit (NEB) for the detection of Mycn and Myc. Primers sequences are specified in Table S3. 4ng (KLF2), 10ng (N-myc), and 2ng (c-Myc) of template were used for qPCR. All cDNA samples were analyzed in triplicates using Fast SYBR green master mix (Applied Biosystems) for ΔΔCt analysis with QuantStudio 5 Real-Time PCR System, 384 wells thermal cycler (Applied Biosystems). Relative expression levels were calculated as a ratio to the average value of house-keeping genes, Proteasome subunit beta type-8 (PSMB8), and hypoxanthine phosphoribosyltransferase 1 (HPRT1). Results show gene expression normalized to PSMB8.

**In vivo polarization.** To induce unspecific in vivo polarization of $CD4^+$ T cells C57BL/6 mice were subcutaneously injected with two subsequent doses of seasonal influenza vaccine INFLUVAC (Abbott Biotechnologies; 1:20 of the human dose) and IFA (incomplete Freund's adjuvant; Sigma) at 1:1 ratio, on days 1 and 14. Following 48h from the second injection, cells were collected from spleens and lymph nodes. Spleens were harvested and mashed into 70µm cell strainer and red blood cells were lysed using ACK buffer (Lonza, Basel Switzerland). Lymph nodes were harvested from inguinal area, and mashed into 70µm cell strainer. Cells were washed, stained with antibodies, and further analyzed using flow cytometry.

**Antibodies and Cell Staining.** All antibodies and dyes are specified in Table S1 (antibodies) and Table S2 (dyes). *Imaging flow cytometry.* Cells prepared for size measurements were collected and stained with live-dead blue reagent (Invitrogen) for 1 hour at 4°C. Next, cells were stained with anti-CD4 and anti-CD44, and were imaged by an Imaging Flow Cytometer (single camera ImageStreamX, AMNIS corp. - part of Luminex, TX, USA). Cells prepared for N-Myc expression and localization quantification were collected, washed with staining buffer (PBS+ 1% BSA), and were extracellularly stained with anti-CD4. Next, cells were fixated with 80% Methanol for 5 min at room temperature (RT), and further permeabilized with 0.1% Tween for 20 min at RT. Following permeabilization, cells were intracellularly stained with anti-Nmyc antibody. First, cells were incubated with the primary anti-Nmyc antibody for 30 min at RT. Next, cells were washed and incubated with the secondary goat anti-mouse antibody for 30 min at RT in the dark. Following intracellular staining, the cells were added with Hoechst 33342 (Invitrogen) to identify cells nuclei, and imaged in ImageStreamX mark II (AMNIS corp. - part of Luminex, TX, USA). *In vivo polarization.* Cells were collected, washed with FACS staining buffer (PBS supplemented with 2% fetal bovine serum and 1 mM EDTA) and incubated with Fc receptor blocker (TrueStain fcX; BioLegend) for 5 min at 4°C. To differentiate between live and dead cells, a viability staining step was done using the eFluor780-Fixable Viability Dye (eBioscience) following the manufacturer's instructions. Cells were then washed and incubated with surface antibodies against CD4 and CD44 for 20 min at 4°C, and were washed twice with a FACS staining buffer. Following staining for surface markers, intracellular labeling was performed. Cells were fixated and permeabilized using the Foxp3/ Transcription Factor Staining buffer set (eBioscience), blocked with Rat serum (1μl per 100μl of staining buffer), and stained with the following antibodies: anti-GATA3, anti-T-bet, anti-RORγt, and anti-Foxp3. Samples were measured using flow cytometry (CytoFLEX, Beckman Coulter) and data were further analyzed with FlowJo software (TreeStar) and Matlab.

**Size measurements.** To assess the size of cells four instruments were used: LSRII flow cytometer (BD, forward scatter (FSC) measurements), Eclipse iCyt flow cytometer (Sony Biotechnology, ec800 software; volume measurements), ImageStream (area measurements), and Multisizer IV Coulter Counter (Beckman Coulter, volume measurements).

**Imaging flow cytometry analysis.** *To assess cell size*, cells were imaged by an Imaging Flow Cytometer (single camera ImageStreamX, AMNIS corp. - part of Luminex, TX, USA). Data was acquired using a 60X lens (NA=0.9), and lasers used were 405nm (125mW), 561nm (200mW), and 658nm (120mW). Data was analyzed using the manufacturer's image analysis software IDEAS 6.2 (AMNIS corp.). Images were compensated for spectral overlap using single-stained controls. Single cells were selected by plotting the area (in µm2) vs. the aspect ratio of the Bright-field image. To eliminate out of focus cells, cells were further gated using the Gradient RMS (measures the sharpness quality of an image by detecting large changes of pixel values in the image). Viable cells were selected by plotting the intensity of Live/dead stain (channel 1) vs. the area of the bright-field image (positively stained cells were considered as non-viable). Cells were then gated according the intensity (the sum of the background subtracted pixel values within the masked area of the image) of CD4 (channel 3) and CD44 (Channel 6) for CD4+/CD44+ and CD4+/CD44- populations. *To quantify N-Myc expression and localization*, cells were imaged with a dual camera ImageStreamX mark II (AMNIS corp. - part of Luminex, TX, USA), using a 60X lens (NA=0.9), and lasers used were 405nm (3mW), 488nm (100mW), 561nm (200mW), and 642nm (150mW). Cells were first gated for DNA staining by plotting the intensity vs. area of the Hoechst 33342 staining (Channel 7). Cropped cells were eliminated by plotting the area of the bright-field image vs. the Centroid X feature (the distance of the cell from the left side of the image field). To further eliminate apoptotic cells, the contrast of the bright-field image was plotted vs. the area of the top 50% intensity pixels of the Hoechst 33342 staining (Area_Threshold (M07, Hoechst, 50)). High contrast, low DNA area cells were considered as apoptotic. $CD4^+$ cells were gated using the intensity vs. Max Pixel (value of the highest intensity pixel within the image) features of the PE-CD4 staining (Channel 3). Aberrant or corrupted cells were further eliminated by gating only on cells with membrane CD4 staining, by using the Max Contour Position (the location of the contour in the cell that has the highest intensity concentration, mapped to a number between 0 and 1, with 0 being the object center and 1 being the object perimeter) and the internalization feature (the ratio of the intensity inside the cell to the intensity of the entire cell). High max-contour position and low internalization cells were considered as CD4 membrane staining and were gated for further analysis. The co-localization between N-Myc and the nucleus were calculated using the Similarity feature (log transformed Pearson's Correlation Coefficient and is a measure of the degree to which two images are linearly correlated within a masked region) between cells with positive N-Myc staining (Alexaflour 647, Channel 11) and the Hoechst 33342 staining.

**Data analysis.** RNA sequencing data was mapped to MM9 genome. Number of reads was normalized to 10,000/sample and low-read genes (<=50) were filtered out. Differential expression analysis was performed with DESeq2 R package (Bioconductor) and other R scripts. Unless otherwise stated, differentially expressed genes were considered to have log2-fold-change (LFC) higher than 2, standard deviation > 0.5 and p-value <=0.05. We have used hierarchal clustering with several agglomeration methods, including Centroid-linkage and Complete-linkage clustering. Gene-set enrichment analyses and interaction analysis were performed with FGSEA (63) and STRING (46).

**Hierarchical-additive mathematical model.** We have solved the following linear regression model:

$$Out_i = A_{i,1} \cdot I_1 + A_{i,2} \cdot I_2 + \cdots + A_{i,6} \cdot I_6 + B_i$$

Where $Out_i$ is the measured output data, either FSC or SSC values; $I_j$ $(j=1,\ldots,6)$ equals 1 if input $j$ was added to the culture medium, and 0 otherwise; $A_{i,j}$ are the model coefficients that describe the influence of input $j$ on output $i$ ; and $B_i$ are model coefficients that describe effective contribution to the output that is not directly regulated by the input cytokines, such as the influence of cytokines secreted by the T cells themselves. We solved the set of 64 linear equations per PC dimension, and obtained the best-fit solution of 7 parameters for each dimension ($A_{i,1}$ , ... , $A_{i,6}$ , $B_i$). See (17) for more details.

**Software.** Data analysis was conducted using Matlab (MathWorks, Natick, MA), R Studio (RStudio Inc., Boston, MA), JupyterLab, FCS Express (De Novo Software, Pasadena, CA), FlowJo (TreeStar) and Easyflow (64) – a Matlab package that was developed in our lab to analyze flow cytometry data.

### Supporting tables

**Table S1. Antibodies specification: cell culture, flow cytometry, and imaging flow cytometry**

| Antibody specificity | Format | Clone | Purchased from | Usage |
|---|---|---|---|---|
| **Cell culture** | | | | |
| α-CD3 | Purified | 145-2C11 | Biolegend | Cell culture |
| α-CD28 | Purified | 37.51 | Biolegend | Cell culture |
| **Flow cytometry/ Imaging flow cytometry (Fluorescence)** | | | | |
| α-CD4 | Brilliant Violet 510 | GK1.5 | Biolegend | Surface staining |
| α-CD4 | PE | RM4-5 | Biolegend | Surface |
| α-CD4 | Pacific blue | RM4-5 | Biolegend | Surface |
| α-CD44 | Brilliant Violet 650 | IM7 | Biolegend | Surface |
| α-CD44 | APC-Cy7 | IM7 | Biolegend | Surface |
| α- GATA3 | PerCP Cy5.5 | 16E10A23 | Biolegend | Intracellular staining |
| α-Tbet | Brilliant Violet 421 | 4B10 | Biolegend | Intracellular |
| α-RORγt | PE | AFKJS-9 | eBioscience | Intracellular |
| α- Foxp3 | Alexa Fluor 488 | 150D | Biolegend | Intracellular |
| α-Nmyc | Purified | NCM II 100 | Abcam | Intracellular, primary Ab |
| Goat α-mouse IgG | Alexa Fluor 647 | Polyclonal | Abcam | Secondary Ab |

**Table S2. Fluorescent dyes used for flow cytometry and imaging flow cytometry**

| Dye description | Usage | Purchased from |
|---|---|---|
| CFSE | Proliferation dye | Invitrogen |
| Hoechst 33342 | Nucleic acid stain | Invitrogen |
| Live-dead blue | Viability dye | Invitrogen |
| 7-AAD | Viability dye | Biolegend |
| eFluor-780 | Viability dye | eBioscience |

**Table S3. qPCR primers. Related to Figure 6.**

| Gene | Forward primer | Reverse primer |
|---|---|---|
| *klf2* | CTCAGCGAGCCTATCTTGCC | CACGTTGTTTAGGTCCTCATCC |
| *n-myc* | CCTCCGGAGAGGATACCTTG | TCTCTACGGTGACCACATCG |
| *c-myc* | GCTGGACACGCTGACGAAA | TCTAGGCGAAGCAGCTCTATTT |
| *hprt* | ATTCAGGAGAGAAAGATGTGATTGA | AGCCAACACTGCTGAAACAT |
| *psmb8* | TGGAGAGTTCCGATGTCAGT | ACTGAAGTAGTCCCAGGTCTC |

**Table S4. Gene interaction network references. Related to Figure 5.**

| Node 1 | Node 2 | Type of interaction | Relevant references |
|---|---|---|---|
| Cell activation | Nfil3 | Induction/ activation | (28, 65) |
| Cell activation | Atf3 | Induction/ activation | (66–68) |
| Cell activation | Bmp2 | Induction/ activation | (69, 70) |
| Cell activation | Yap1 | Induction/ activation | (71, 72) |
| Cell activation | Pparg | Induction/ activation | (73–75) |
| Cell activation | Mycn | Induction/ activation | (76) |
| Nfil3 | Smad7 | Induction/ activation | (28) |
| Ahr | Atf3 | Induction/ activation | (77–79) |
| Bmp2 | Pparg | Induction/ activation | (80, 81) |
| Yap1 | Bmp2 | Induction/ activation | (82) |
| Yap1 | Pparg | Inhibition/ repression | (83, 84) |
| Yap1 | Smad2/3 | Complex crosstalk | (85–88) |
| Smad6 | Bmp2 | Inhibition/ repression | (89–91) |
| Smad7 | Yap1 | Induction/ activation | (92, 93) |
| Smad7 | Smad2/3 | Inhibition/ repression | (94–97) |
| Smad2/3 | Nfil3 | Inhibition/ repression | (28, 65, 98) |

| | | | |
|---|---|---|---|
| Smad2/3 | Atf3 | Induction/ activation | (90, 99–101) |
| Smad2/3 | Pparg | Inhibition/ repression | (102–105) |
| Smad2/3 | Ahr | Induction/ activation | (106, 107) |
| Pparg | Smad2/3 | Inhibition/ repression | (80, 108–110) |
| Tgfbr1 | Smad2/3 | Induction/ activation | (111–113) |
| Smad2/3 | Smad7 | Induction/ activation | (94–96, 114, 115) |
| Tgfbr1 | Mycn | Inhibition/ repression | (51, 52, 60, 61) |
| Tgfbr1 | Klf2 | Induction/ activation | (116, 117) |
| Ahr | Pparg | Inhibition/ repression | (118, 119) |
| Ahr | Mycn | Inhibition/ repression | (120) |
| Ahr | Cell growth | Complex crosstalk | (121–123) |
| Klf2 | Smad7 | Induction/ activation | (116, 124) |
| Klf2 | Pparg | Inhibition/ repression | (125–127) |
| Klf2 | Cell growth | Inhibition/ repression | (47, 125, 128) |
| Mycn | Klf2 | Induction/ activation | (129) |
| Mycn | Cell growth | Induction/ activation | (48, 56, 57, 130–133) |

### Supporting figures

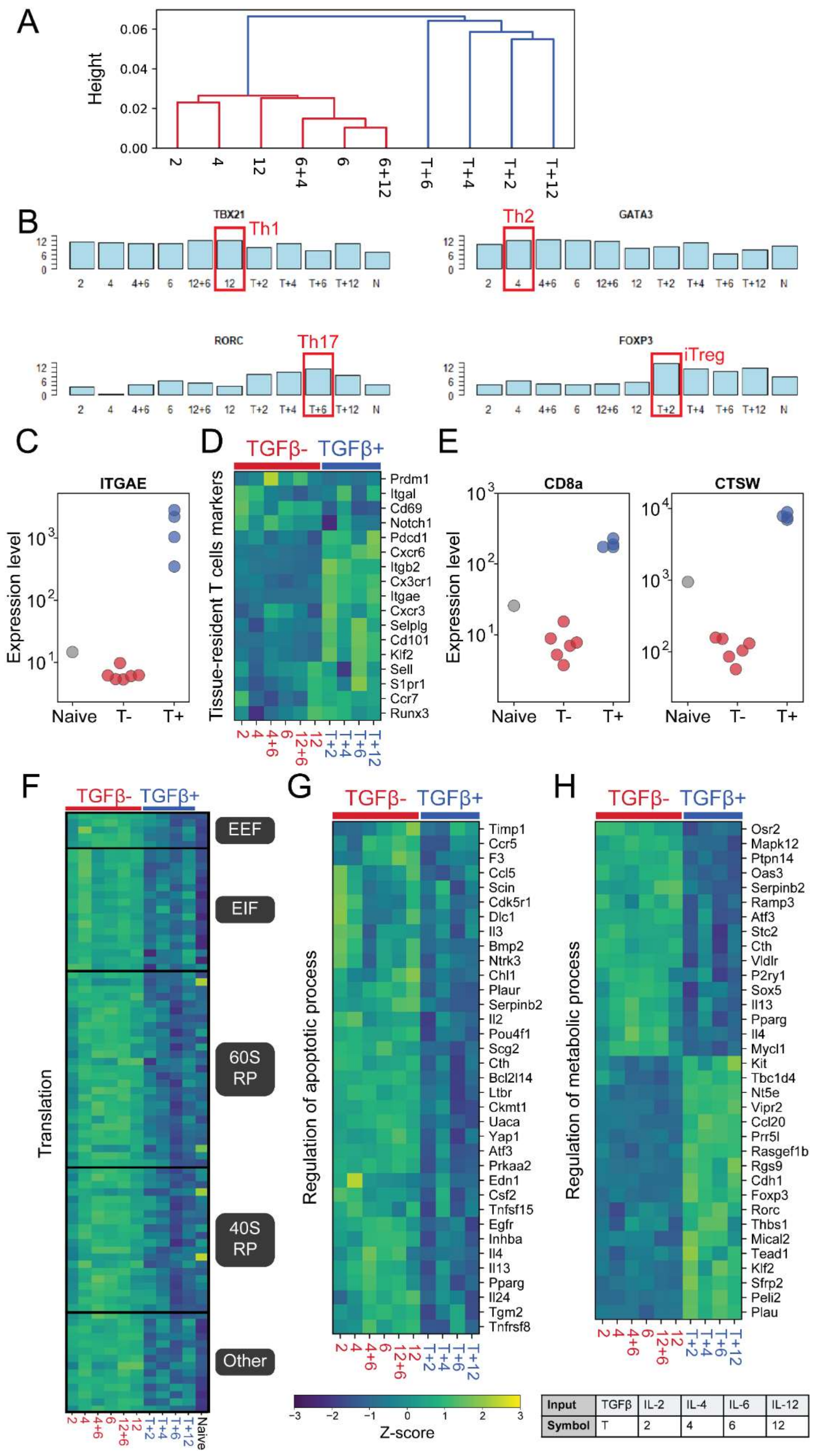

**Figure S1. TGFβ, in combination with other cytokines, induces changes in genes associated with CD4⁺ T-cell differentiation, tissue residency, translation, cell death, and metabolism. Related to Figure 1 and Figure 2.** Naïve $CD4^+$ T cells were cultured under different cytokine conditions, in the presence and absence of TGFβ. Cells were harvested after 96h of culture, and gene expression was measured using RNA-seq. Cytokine combinations are marked using one-symbol labels, as specified in the table at the bottom. Results are averaged over n = 3 repeats. (A) Hierarchical clustering was performed on the sequencing data. Sample are clustered according to the correlation between their gene expression profile. Distance was calculated to be the Spearman Correlation coefficient between each two conditions. (B-H) Normalized expression levels of: (B) Lineage-specifying transcription factor expression under the different input conditions. We examined the expression of the TFs characterizing the four main Th fates: TBX21 (Th1, top-left), GATA3 (Th2, top-right), RORc (Th17, bottom-left), and FOXP3 (Treg, bottom-right). Classical Th-subset inducing conditions are marked by red boxes; N – Naïve cells. (C) Expression of ITGAE, the gene encoding the tissue-resident memory T cells marker CD103, in the presence or absence of TGFβ. (D) Expression of genes that are involved in the differentiation and maintenance of tissue-resident memory T cells, in the presence or absence of TGFβ ($SD \geq 0.5$; $P$-value$\leq 0.05$, see Methods). (E) Expression of the cytotoxic T cells markers, CD8a and CTSW, in the presence or absence of TGFβ. (F) Expression level heat map of the gene-set enrichment analysis (GSEA) leading-edge subset of the "Translation" gene set. Data is shown for activated cells cultured under different cytokines conditions and compared to naïve cells. (G) Genes involved in the regulation of apoptotic processes, in the presence or absence of TGFβ ($SD \geq 0.5$; P-value$\leq 0.05$). (H) Genes involved in the regulation of metabolic processes, in the presence or absence of TGFβ ($SD \geq 0.5$; P-value$\leq 0.05$).

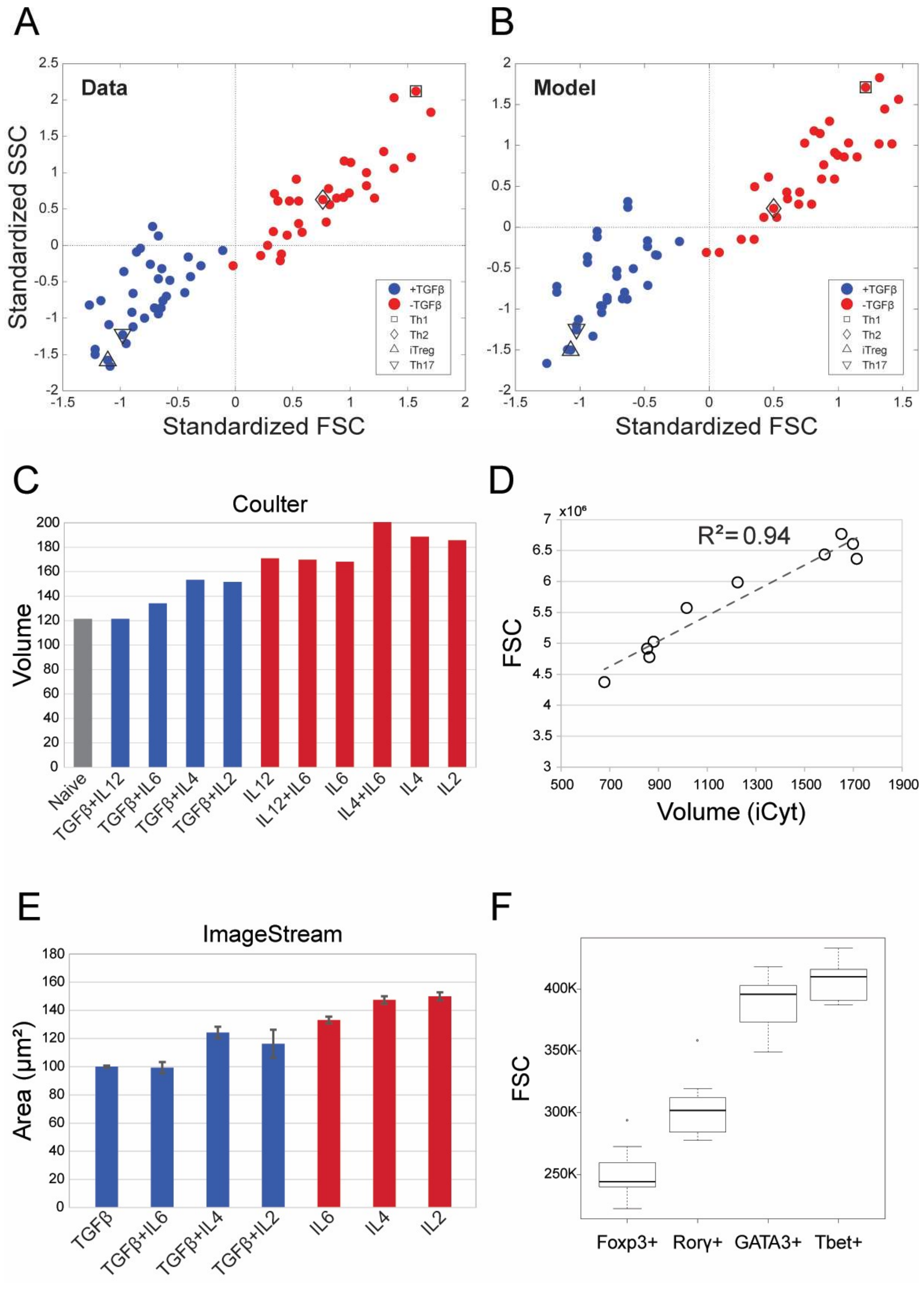


**Figure S2. TGFβ affects the size of in-vitro and in-vivo activated CD4⁺ T-cells. Related to Figure 3.**

(A) Mean FSC and SSC values for 64 different cytokine combinations are plotted. Values were standardized to have zero mean and unit standard deviation. Samples are colored according to the presence (blue) or absence (red) of TGFβ in the culture medium. Conditions that drive polarized Th lineages are marked. The mean values of three independent experiments were used in the analysis. (B) A hierarchical-additive mathematical model (see Methods) that we have previously developed predicts and reproduces the pattern of FSC-SSC values across all conditions. (C) The mean volume of cells cultured under different conditions with (blue) and without (red) TGFβ, as measured using coulter counter. (D) Correlation between volume of cells, measured using iCyt, and forward-scatter (FSC), measured using conventional flow cytometry, demonstrates that FSC constitutes a reliable parameter for relative cell size. (E) Area measurements of cells cultured under different conditions with (blue) and without (red) TGFβ, as measured using imaging flow cytometry (ImageStreamX). The mean and standard deviation (error bars) of four experiments are shown. (F) $CD4^+$ T-cell size in vivo. C57BL mice were injected with influenza+ adjuvant on days 1 and 14. 48h following the second immunization, $CD4^+$ T cells were isolated from spleens (Fig. 3E, main text) and lymph nodes. Cells were stained with antibodies for Th subset-characterizing transcription factors, and the size and phenotype of the cells were examined using flow cytometry. FSC was used as a size indicator. Cells were gated according to the expression of the TFs, which are associated with the four main Th subsets: T-bet (Th1), GATA3 (Th2), RORγt (Th17), and Foxp3 (Treg). Cells that require the presence of TGFβ for their differentiation (Th17, Treg) display a reduced size compared to cells that do not require the presence of TGFβ to induce their differentiation (Th1, Th2). Results are from n=7 mice.

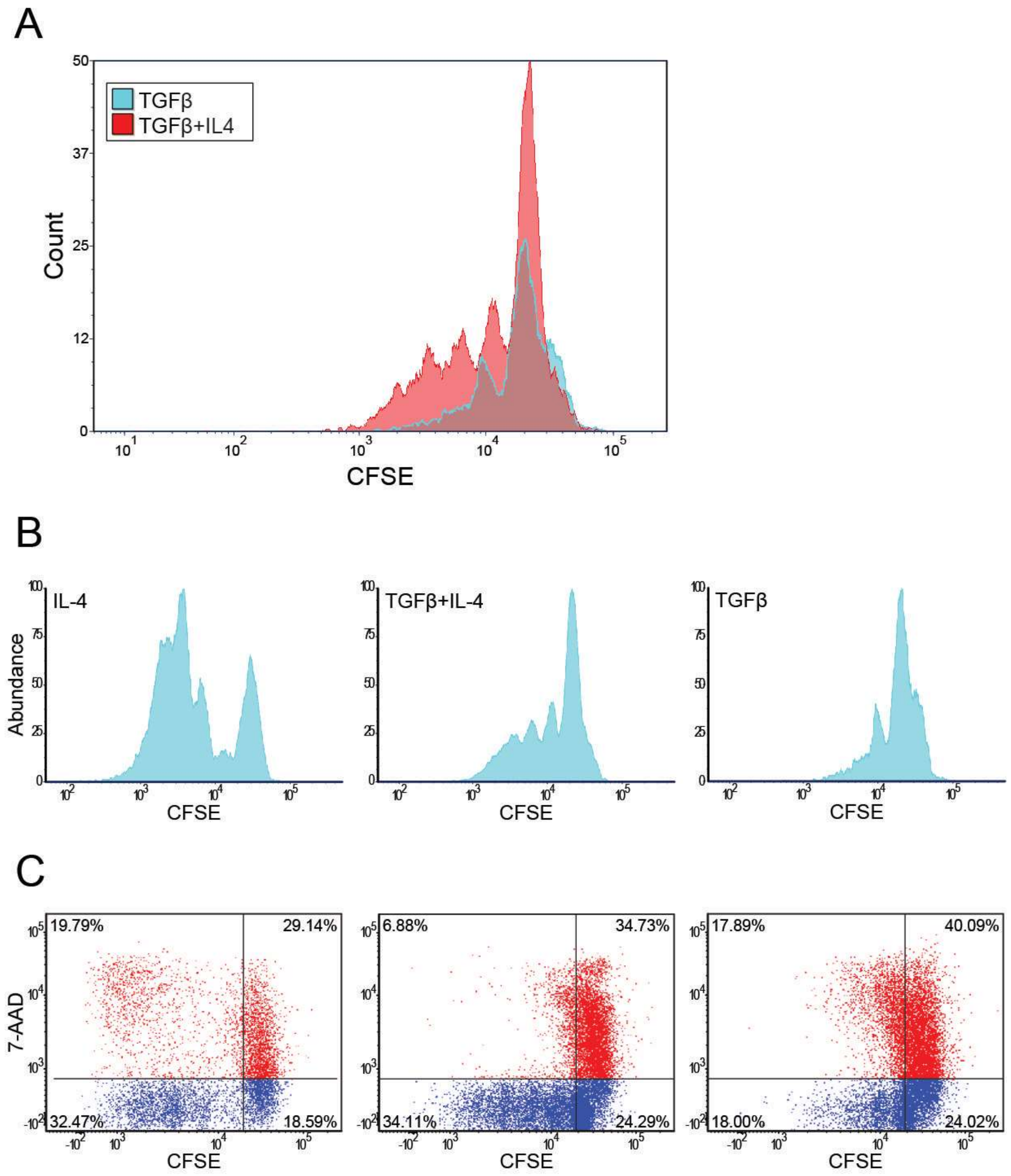


**Figure S3. TGFβ influences the proliferation and survival of CD4⁺ T cells. Related to Figure 4.** Naïve CD4⁺ T cells were isolated, stained with the proliferation dye CFSE, and cultured under different cytokine combinations, as indicated. Proliferation and death of the cells were examined in intervals of 24h. (A) The proliferation of cells cultured with TGFβ alone (light blue) and cells cultured with TGFβ together with an additional cytokine (TGFβ+IL-4, red). (B) Proliferation profiles of three representative conditions: IL-4 (left), TGFβ+ IL-4 (middle), and TGFβ (right) following 96h of culture. (C) Scatter plots showing cell death (by 7-AAD staining) and proliferation (by CFSE), following 72h of culture under three representative conditions: IL-4 (left), TGFβ+IL-4 (middle), and TGFβ (right).
